# Spectral decomposition unlocks ascidian morphogenesis

**DOI:** 10.1101/2023.08.22.554368

**Authors:** Joel Dokmegang, Emmanuel Faure, Patrick Lemaire, Ed Munro, Madhav Mani

**Author notes:** **For correspondence:** (NU); (NU).

## Abstract

Describing morphogenesis generally consists in aggregating the multiple high resolution spatiotemporal processes involved into reproducible low dimensional morphological processes consistent across individuals of the same species or group. In order to achieve this goal, biologists often have to submit movies issued from live imaging of developing embryos either to a qualitative analysis or to basic statistical analysis. These approaches, however, present noticeable drawbacks, as they can be time consuming, hence unfit for scale, and often lack standardisation and a firm foundation. In this work, we leverage the power of a continuum mechanics approach and flexibility of spectral decompositions to propose a standardised framework for automatic detection and timing of morphological processes. First, we quantify whole-embryo scale shape changes in developing ascidian embryos by statistically estimating the strain-rate tensor field of its time-evolving surface without the requirement of cellular segmentation and tracking. We then apply to this data spectral decomposition in space using spherical harmonics and in time using wavelets transforms. These transformations result in the identification of the principal dynamical modes of ascidian embryogenesis and the automatic unveiling of its blueprint in the form of scalograms that tell the story of development in ascidian embryos.

## Introduction

Morphogenesis, the emergence of shape in living systems, is a continuous process littered with spatiotemporal dynamics at various timescales and lengthscales and significance. Developmental biology aims at the identification, localisation, and timing of these processes. Once this work is carried out in a given species, embryogenesis can then be described as a series of stages delineated in space and time by the identified landmarks ***Satoh (1978); Nishida (1986); Jeffery (1992); Keller et al. (2003); Lemaire (2009); Sherrard et al. (2010); Hashimoto et al. (2015); Hashimoto and Munro (2018); Guignard et al. (2020***). In order to rigorously define development landmarks, biologists have mostly had to submit imaged embryos either to qualitative analyses, or to rudimentary statistical analysis. These methods however present major drawbacks. On the one hand, they can be time consuming, hence unfit for scale. On the other hand, since morphogenetic processes tend to be unique to a species, these simple methods often lack a general language and framework that permit comparative analyses. For instance, whereas the analysis of cell counts can inform about the proliferation dynamics in a tissue, it does not reveal anything about the shape of the system. For this purpose, other measurements such as length, width, height, aspect ratios or curvatures would be more suitable. Although efforts have been made to automate the staging of development in living systems ***Jones et al. (2022***), these methods still rely on preliminary examination using traditional methods.

A standardized method able to identify key milestones in development and lay out the blueprint of morphogenesis in a given system is henceforth needed. Recent breakthroughs in microscopy technology have propelled the resolution of live imaging data to the sub-cellular scale, allowing for the uncovering of precise cell and tissue shape dynamics ***Tassy et al. (2006); Stelzer (2015); Power and Huisken (2017***). These advances have created an unprecedented opportunity for the leveraging of computational methods in the study of morphogenesis ***Tassy et al. (2006); Michelin et al. (2015); Stegmaier et al. (2016); Leggio et al. (2019); Guignard et al. (2020***). The rigorous and physically motivated framework of continuum mechanics accommodates itself well to the flow-like dynamics of biological tissues ***Humphrey (2003); Ambrosi et al. (2011); Blanchard et al. (2009); Humphrey (2013); Streichan et al. (2018***). Within this framework, strain-rate fields, which measure the rate at which the shape of a system changes with time, are suited to characterise the dynamical behaviour of the system. Moreover, mounting evidence have informed of the requirement for embryo-wide approaches in the study of morphogenetic flows ***Streichan et al. (2018); Mitchell et al. (2022***). However, although the evaluation of such global fields across the spatial and temporal domains spanned by a system of interest may reveal valuable insights into its dynamical workings ***Bar-Kochba et al. (2015); Stout et al. (2016); Patel et al. (2018***), their sole determination might not be sufficient for a holistic description of the behaviour of the system: there is a need for novel methods to analyse them.

This is especially true when it comes to morphogenesis ***Dalmasso et al. (2021); Romeo et al. (2021); Mitchell and Cislo (2022***). The processes involved in development are inherently multiscale, both in the spatial and temporal domains, and may interact or overlap ***Godard and Heisenberg (2019); Dokmegang et al. (2021); Dokmegang (2022***). As it is the case with several species ***Godard and Heisenberg (2019***), ascidian early development is a playground featuring important displays of cellular divisions and tissue mechanics ***Lemaire (2009***). The local behaviors captured by indicators such as the strain-rate field might therefore arise from a non-trivial superposition of these dynamical modes, essentially making these measurements complex to interpret without further analysis. Spectral decomposition, whereby a signal is broken down into its canonical components, is well suited to the study of systems that exhibit multimodal behaviors ***Romeo et al. (2021); Dalmasso et al. (2021***). The benefits are at least two-folds: (i) individual constituents may represent distinct dynamical processes, thereby enabling the decoupling of physical processes entangled in the data; (ii) only a handful of components may significantly contribute to the original function, resulting in a compressed, lower dimensional representation that capture the main features of the studied process. The canonical components usually take the form of well known families of functions whose linear combination can reconstitute the original field.

In this work, we take advantage of microscopy imaging data to develop a generic computational framework able to identify and delineate the main features of morphogenesis. Our method takes as input *3D+time* images of developing ascidian embryos and outputs spatiotemporal scalograms of ascidian development in the form of heatmaps that highlight key developmental processes and stages of ascidian gastrulation. By virtue of a novel meshing scheme derived from level-set methods, raw cell geometry data is first transformed into a single time-evolving embryonic surface on which the strain-rate tensor field can be computed. The accuracy of our inference of a strain-rate field relies on high-frequency temporal sampling, characterised by small deformations of the embryonic surface between subsequent time points. The morphomaps we present are a result of spectral analyses of the strain-rate fields, featuring spherical harmonics decomposition in the spatial domain and wavelet decomposition in the temporal domain. In summary, our method can identify and classify dynamical morphogenetic events. In particular, we are able to identify and distinguish the morphogenetic modes of gastrulation and neurulation phases in ascidian development, recover the characteristic two step sequence of endoderm invagination originally described using 3D analysis of cell shapes ***Sherrard et al. (2010***), and capture patterns of cellular divisions in ascidian development ***Nishida (1986***). Moreover, our method identifies a distinctive stage of ascidian gastrulation, *‘blastophore closure’*, which follows endoderm invagination and precedes neurulation.

## Results

### Definition of Lagrangian markers on the surface of the embryo

In order to recover a continuum description of the dynamics in ascidian morphogenesis, we aim to examine the time evolution of strain-rate fields across the entire surface of developing embryos. This endeavour however presents at least two significant challenges. On the one hand, strain-rate computation requires the presence of fiducial markers on the surface of the embryo. Characteristically, this requirement is not always accounted for in the imaging of developing embryos. On the other hand, the outer layer of the embryo being constituted of single cell apical faces, even if such markers had been defined at an earlier time point, uncontrolled stochastic biological processes such as proliferation within the tissue might subsequently grossly uneven the distribution of these markers, thus rendering the computed mechanical indicators at best imprecise. Given the non-triviality of an experimental setup able to solve the described issues, a computational method is required. The goal of such a method would be to computationally discretise an embryo surface into a set of material particles whose trajectories can be tracked in small lapses of development. The positions of these markers over time can then be used to derive mechanical indicators of development dynamics ***Humphrey (2003***).

To achieve this fit, we first take advantage of the level set scheme described in ***Zhao et al. (2000***) to define static markers on the surface of the embryo at every timepoint of development. The gist of our method resides in the definition of a homeomorphic map between the surface of the embryo (*S*_1_(*t*)) and a topologically equivalent mesh (*S*_2_(*t*)) whose number of vertices, faces and edges remain fixed (fig. 1a). Conformal parametrisations of embryonic shape have been used in other systems ***Alba et al. (2021***). Here, the topologically equivalent mesh consists of a sphere resulting from successive butterfly subdivisions of an icosahedron ***Hardy and Steeb (2008***). Using a homeomorphic map, this mesh can be deformed to match the surface of the embryo at each time point of development (fig. 1b). As in ***Zhao et al. (2000***), the map is obtained by finding the positions of *S*_2_(*t*) vertices that minimize the distance between both surfaces (fig. 1a, *right*). At the initial time point, *S*_2_(*t*) is chosen to be a sphere enclosing the embryo. For various reasons, including computational efficiency, variants of this method can be defined such that, for instance, at subsequent time steps, linear combinations of the sphere and its deformations matching the embryo at preceding steps are used. Further details are given in the supplementary materials.

**Figure 1.**
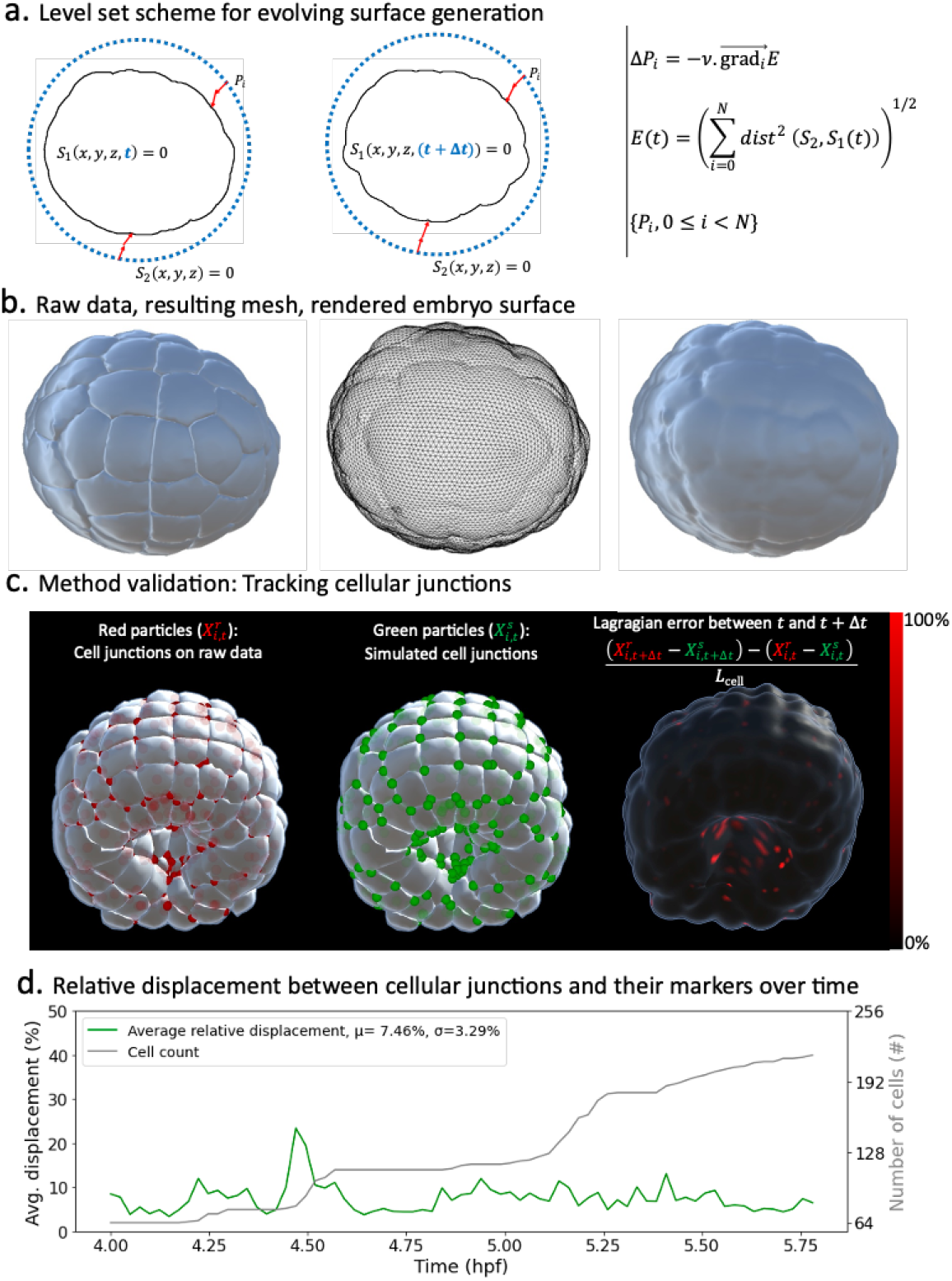
Level-sets inspired Lagragian markers. **a)** *Left* Schematics of the level set method. *Right* Fundamentals of the numerical scheme that shapes *S*_2_ into *S*_1_. **b)** Illustration of the method in action. *Left* Raw data consisting of geometric meshes of single cells spatially organised into the embryo. *Center* Embryo surface mesh resulting from the application of the level set scheme. *Right* Rendering of the embryonic surface. **c)** Tracking of cellular junctions. *Left* Identification of cellular junctions (*red dots*). *Center* Corresponding markers (*green dots*, defined as vertices on the computed embryonic surface closest to the junctions. *Right* Relative displacement between junctions and their markers at consecutive timepoints. **d)** Plot over time of the relative displacement between cellular junctions and their markers.

Next, via a numerical study, we assert that markers defined as such behave as Lagrangian particles in small increments of developmental dynamics. To support this point, we identify on the raw dataset the positions of cellular junctions at the surface of the embryo and evaluate how well our virtual markers mimic their movements in time. We measure the relative displacement between a cellular junction (fig. 1c *red points*) and its corresponding vertex (fig. 1c, *green points*) at consecutive time points. We take the difference between these distances and normalise it by the average side length of cell apices. Despite gross approximations inherent to the nature of the dataset (geometric meshes) and the process of identifying cellular junctions (averaging the barycenters of closest triangles between three or more cells in contact), the relative displacement between cellular junctions and their markers remains on average relatively small (under 8%, fig. 1d). We further show that this characteristic is independent of the number of particles used in the method (sup fig. 1c), making it a remarkable property of the scheme. This result sheds even more favorable light on the method when considering that cellular junctions, precisely because they are the meeting point of three or more cells, are expected to exhibit more chaotic behaviour than single cell particles. Moreover, these numbers are skewed by large scale morphogenesis processes such as synchronised cell divisions, as evidenced by spikes in fig. 1d, and fast-paced endoderm gastrulation, as highlighted by higher errors at the vegetal pole of the embryo during this phase (red dots in fig. 1c, *right*).

### Strain rate Field describes ascidian morphogenesis

Once a mesh representing the surface has been constructed for the embryo surface, we proceed with the computation of the strain-rate fields across the surface of the embryo and throughout development timeline. Thanks to the Lagrangian nature of mesh vertices, a velocity field can be defined on the mesh. Although particles at every given time point live on the 2D surface of the embryo, their trajectories in time involve greater degrees of freedom in the 3D space. A correct parametrisation of the velocity field at every position henceforth requires three coordinates **v(x)** = (*v*_*x*_(**x**), *v*_*y*_(**x**), *v*_*z*_(**x**))^*T*^. The strain rate field is derived as the symmetric part of the discrete gradient of the velocity field, computed as described in ***Mancinelli et al. (2018***). Intuitively, the strain rate evaluated on a given mesh vertex measures how the velocity vector varies in the neighbourhood this point ***Mancinelli et al. (2018); De Goes et al. (2020***).

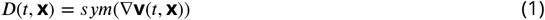

The mathematical construction of the strain rate eq. 1 implies that its algebraic representation takes the form of a second order tensor which can be written down as a 3 × 3 matrix (*D*)_*ij*_. The diagonal elements of this matrix capture the linear strain rate in the *x, y, z* axes, depicting the change in length per unit time. The non-diagonal elements stand for shear strain rates in the the *xy, xz* and *yz* directions. Because *D* is symmetric, there exists an alternative representation which holds stronger local geometric meaning. This representation is obtained by computing the eigenvectors and eigenvalues of the strain rate tensor. Eigenvectors stand for orthogonal spatial directions that are not rotated, but only stretched, by the application of the strain rate matrix. They define the principal axes of a coordinate system in which the strain rate tensor would be solely composed of maximal linear strain rates (fig. 2a). From this decomposition, we derive a scalar field that is computed at every mesh particle as the square root of the sum of the squared eigenvalues of the strain-rate (fig. 2b). Intuitively, this field describes the magnitude of the rate of change underwent by a particle at the surface of the embryo in the three orthogonal spatial directions of most significant rate of change.

**Figure 2.**
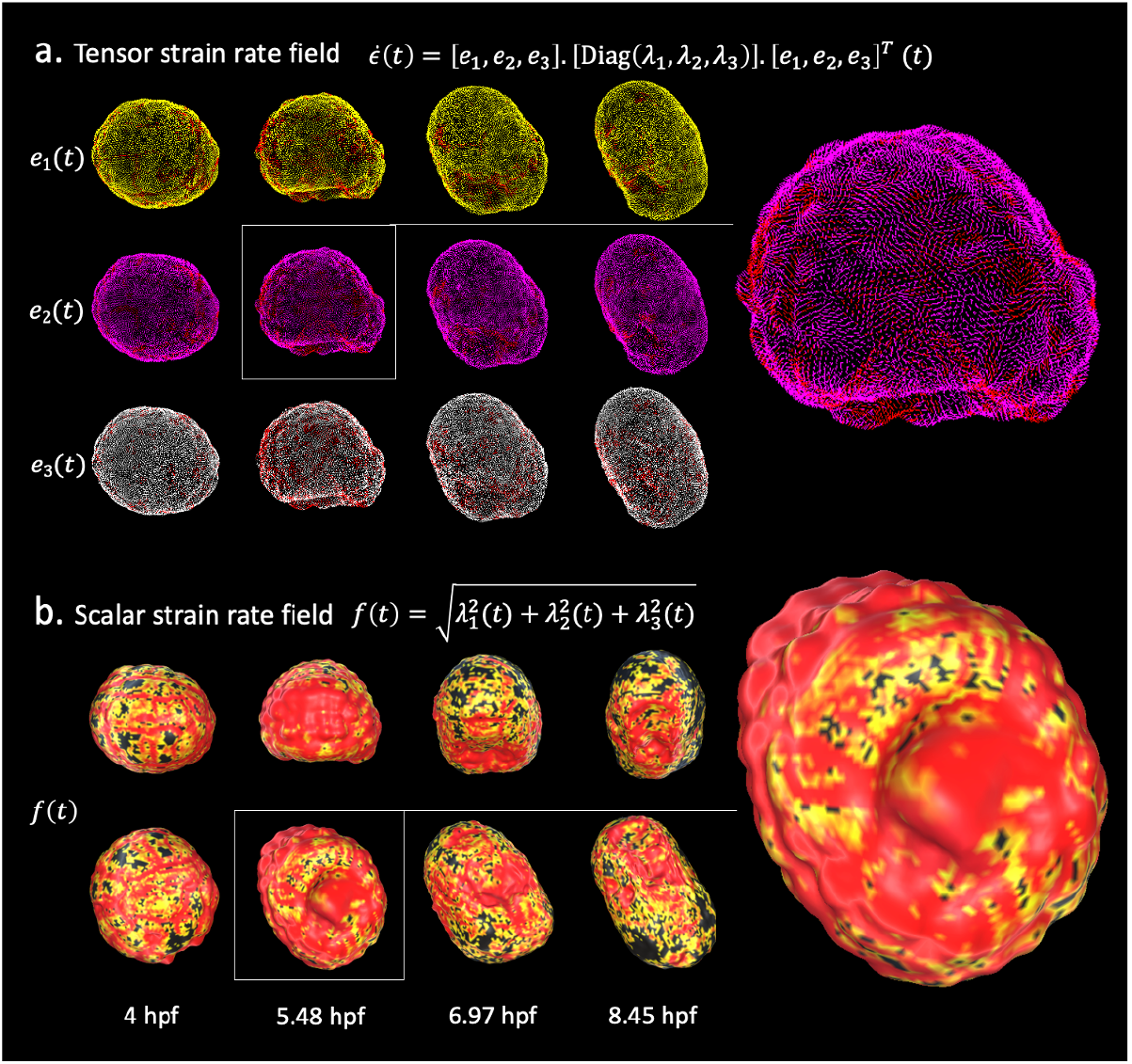
Strain–rate field describes morphogenesis. The strain-rate tensor field measures the rate at which morphological changes occur in the embryo as a function of time. The strain-rate tensor field is locally represented as a 3×3 symmetric matrix and is completely determined by its eigenvector fields. **a)** Heatmap of the eigenvector fields of the strain rate tensor. Each row represents a vector field distinguished by a distinct root color (*yellow, pink, white*). The gradient from the root color to red represents increasing magnitudes of the strain rate tensor. *Top* Spatiotemporal dynamics of the first eigenvector field. *Middle* Spatiotemporal dynamics of the second eigenvector field. *Bottom* Spatiotemporal dynamics of the third eigenvector field. **b)** Heatmap of the scalar strain rate field. The gradient from yellow to red depicts regions of increasing morphological activity, while black stands for areas of low morphological activity. The heatmaps show high morphological activity in the invaginating endoderm and zippering neural plate, but also across the embryonic animal during rounds of synchronized division.

In order to minimize undesirable artifacts that may arise from numerical inefficiencies, we apply a Gaussian filter to the strain rate tensor field before deriving the scalar field. At each particle location, we apply a Gaussian convolution mask spanning its first and second order neighbourhood. A similar smoothing process is also used in the time domain. Interestingly, this strain-rate derived scalar field remarkably mirrors well-known features of ascidian development. Similarities between the spatiotemporal distribution of morphogenesis processes described in the literature and heatmaps of this field on the evolving embryo surface emerge. On the one hand, wider spatial gradients of yellow to red depicting higher morphological activity portray the spatiotemporal locations of endoderm invagination in the embryonic vegetal pole (fig. 2b, *center-left*) ***Sherrard et al. (2010***), synchronised rounds of division in the animal pole, and zippering in the neural plate ***Hashimoto et al. (2015***) (fig. 2b, *center-right, right*). On the other hand, known spatiotemporal locations of low morphological significance (e.g. the animal pole when not proliferating) in the embryo exhibit stronger concentration of mechanical activity on cell boundaries, with the corollary that cellular identities are mostly preserved (fig. 2b, *t* = 4*hpb*). A notable by-product of this scalar field is the evidencing of the duality of the embryo as both a sum of parts constituted of cells and an emerging entity in itself: the strain rate field clearly discriminates between spatiotemporal locations where isolated single cell behaviours are preponderant (e.g. fig. 2b, *t* = 4*hpb*) and those where coordinated cell behaviours dominate (e.g. fig. 2b, *t* = 5.48*hpb*).

This brief overview already demonstrates the riches in a quantitative, spatially global and not event-driven approach to study morphogenesis. It also sets the stage for further analysis of morphogenesis dynamics in the ascidian embryo.

### Spectral decomposition in space: Spherical harmonics reveal the main modes of ascidian morphogenesis

In order to capture relevant features of the strain rate field in the spatial domain, we conduct a spectral analysis of the scalar strain rate field. The family of spherical harmonic functions stands out as a de-facto standard for the study of signals defined on a unit sphere, and by extension on surfaces homeomorphic to the sphere. Spherical harmonics form an infinite orthonormal basis of functions defined on the surface of the sphere and represent a generalisation of the Fourier series for functions of two variables ***Knaack and Stenflo (2005***). Unsurprisingly, these functions play an important role in many branches of science displaying spherical symmetry, including quantum mechanics and geophysics ***Knaack and Stenflo (2005); Dahlen and Tromp (2021***). Spherical harmonics have recently been used in studies of morphogenesis in zebrafish and mouse ***Romeo et al. (2021); Dalmasso et al. (2021***).

Spherical harmonic basis functions are indexed by two parameters (*l, m*), such that *l* ≥ 0, |*m*| ≤ *l* representing respectively the degree and order of the harmonic (Supplementary figure 3). A signal defined on the sphere can be written as a linear combination of such functions. Decomposing a signal into spherical harmonics hence amounts to finding the coefficients *f*_*lm*_ of this weighted sum. In the case of our spatiotemporal scalar strain rate field, the coefficients *f*_*lm*_ are also a function of time and can be obtained as shown in equation 2.

**Figure 3.**
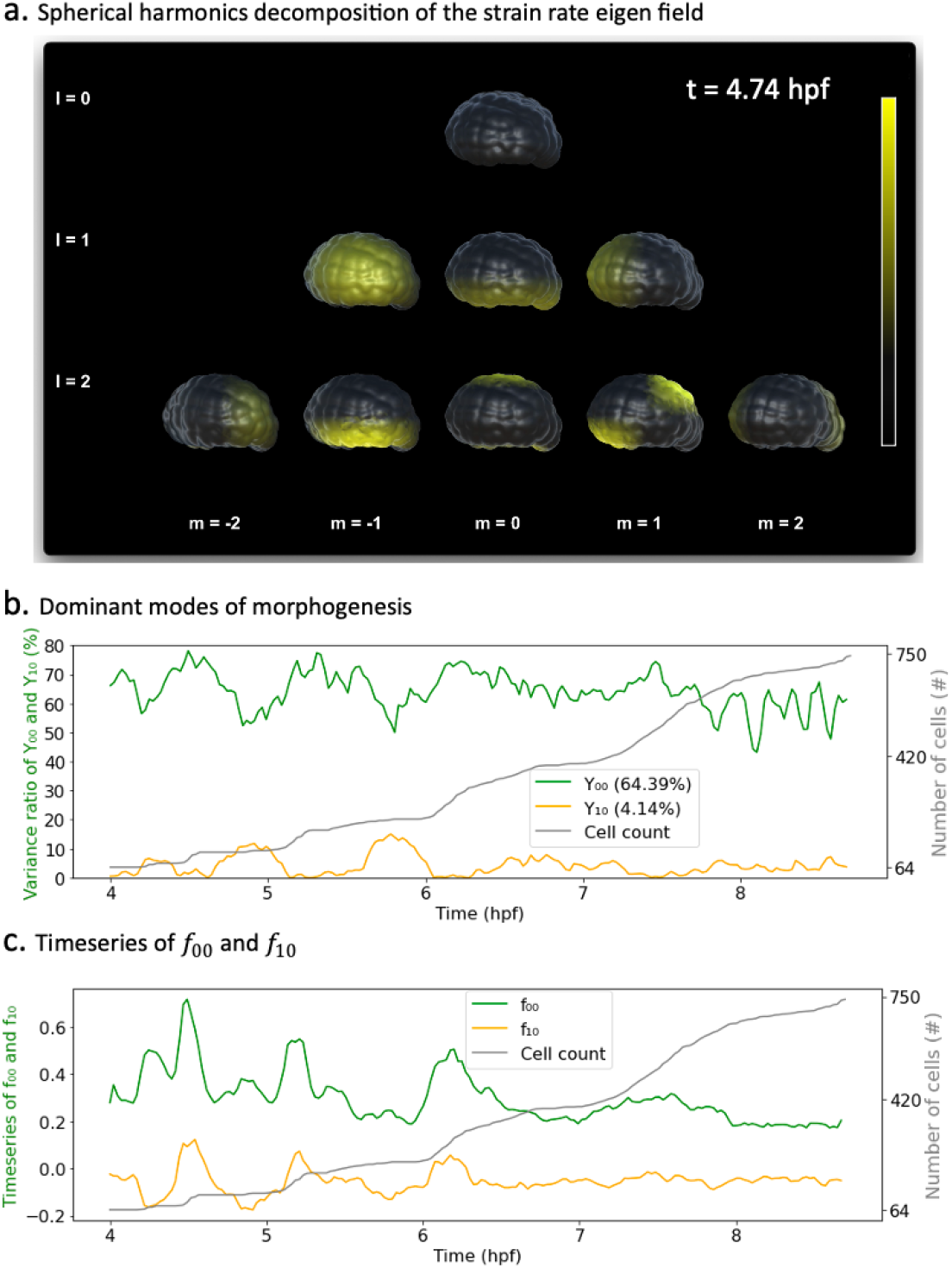
Spherical harmonics decomposition of morphogenesis. **a)** Example of Spherical harmonics decomposition of the scalar strain rate field mapped to the embryo at *t* = 4.74 hpf. Each picture represents the value of the harmonic field (*Y*_*lm*_) multiplied by its coefficient *f*_*lm*_. *f*_*lm*_*s* here are taken relative to each specific *f*_*lm*_ minimal and maximum bounds in the entire time window of observation. Thresholding is applied for better rendering. **b)** Time evolution of the variance ratios of the main modes of ascidian early morphogenesis (*Y*_00_ and *Y*_10_). The cell population dynamic is also included in the plot for clarity. **c)** Time evolution of the coefficients *f*_00_ and *f*_10_ associated with spherical harmonics (*Y*_00_ and *Y*_10_). The cell population count is also included in the plot for clarity.

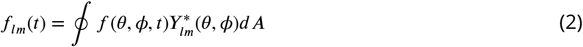

Here, 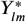 stands for the complex conjugate of the spherical harmonic *Y*_*lm*_. Moreover, for a given degree *l*, each of the (2*l* + 1) spherical harmonics (*Y*_*lm*_) |_*m*_| _<*l*_ spatially partitions the unit sphere into as many spatial domains indicating when a signal is positive, negative or null (Supp. Fig.4a). Figure 3a illustrates the projections of the scalar strain rate field to spherical harmonics (*Y*_*lm*_)_*l*≤2,_ |_*m*_| _<*l*_ at *t* = 4.74 hpf, and their mapping unto the surface of the embryo. These plots reveal for instance that while there is no embryo-wide dominant morphogenesis process at this time (*l* = 0, *m* = 0), smaller regions, notably the vegetal pole are experiencing significant morphological activity (*l* = 1, *m* = 0).

The contributions of each spherical harmonic to the global signal can be assessed more rigorously, and interpreted in the light of biology. To this effect, we observe the temporal dynamics of the coefficients *f*_*lm*_(*t*) associated with each spherical harmonic. In analogy to *Principal Components Analysis*, we measure the average variance ratio 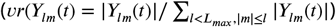 over time of each harmonic with respect to the original signal (Fig. 3b). With an average variance ratio of 64.4%, the spherical harmonic *Y*_00_, capturing embryo-wide morphological activity, contributes the most to ascidian morphogenesis. Spherical harmonic *Y*_10_ is the next contributor, coming second with a variance ratio of 4.1%. This observation is warranted, as *Y*_10_ maps to the animal and vegetal poles of the embryo, which are the epicenters of synchronised cellular divisions and endoderm invagination respectively ***Jeffery (1992); Lemaire (2009***). Interestingly, variances in the directions of *Y*_00_ and *Y*_10_ evolve in an antiphased pattern, most notably in earlier parts of the plot, with *Y*_00_ contributing maximally (and *Y*_10_ minimally), during periods of cell division, before relinquishing some variance shares to *Y*_10_, which then peaks. This suggests that while sporadic deformations induced by cellular divisions often dominate the landscape of morphological activity, an observation consistent with studies in other species ***Cislo et al. (2023***), other localised, slower processes are at play in the embryo. The described pattern tends to fade out in the later parts of the plot, suggesting a shift in development dynamics.

Furthermore, by observing the time dynamics of the coefficients themselves (Fig. 3c), one can easily identify which parts of the embryo are concerned by the morphological changes depicted. For instance, the positive peaks in *f*_10_(*t*) (Fig. 3c) indicate that the morphological processes at hand take place in the northern hemisphere of the sphere. Remarkably, these coincide with rapid growth in cell population and thus synchronous cell divisions, which are known to be restricted to the animal pole of the ascidian embryo during endoderm invagination ***Jeffery (1992***). In addition, most of the dynamics captured by *f*_10_(*t*) are in the negative spectrum (*f*_10_(*t*) < 0), pointing to the lower hemisphere of the embryo, the foyer of several cell shape deformations at play in ascidian early development.

The sporadic short-time scale cell division events in the animal pole coexist with numerous other features of morphogenesis, most notably, the larger scale continuous deformation process in endoderm invagination at the embryonic vegetal pole. Beside the peaks on the plots of the time series, it is not a trivial task to identify what other rich insights may be hidden in this data. A simple observation of the oscillatory patterns of these main modes hence paints an incomplete picture of ascidian morphogenesis. Extracting the footprint of all morphogenesis processes in these time series requires further analysis.

### Spectral decomposition in time: Wavelets analysis of spherical harmonic signals unveils the blueprint of morphogenesis

Analysing timeseries often implies the understanding of how a signal is composed and how its components overlap in time. Wavelets have been put forward as effective multi-resolution tools able to strike the right balance between resolution in time and resolution in frequency ***Torrence and Compo (1998); Lau and Weng (1995***). Although they have been taken advantage of in the broader context of biology, most notably in the analysis of brain and heart signals ***Brunton and Kutz (2022***), they have so far been under-used in developmental biology. The reason might be found in the reality that morphogenesis data is often not understood in term of time series. Our spherical harmonics decomposition of morphogenesis, inspired by similar endeavours in other fields ***Dahlen and Tromp (2021); Knaack and Stenflo (2005***), offers an unprecedented opportunity to leverage the existing rich signal processing toolbox in development biology. In particular, enlisting the help of wavelet transforms in unlocking the complex entanglements of the multiple morphological process at play during ascidian early development. We proceed to apply the Ricker wavelet transform to our spherical harmonics time series, normalised by mean and standard deviation in different time windows of interest. The result is a set of scalograms which decompose the signals into canonical components organised in timelines that reveal the story ascidian morphogenesis.

First, we apply the wavelet transform on the timeseries *f*_00_(*t*) to the entire time range covered by our dataset, comprising both gastrulation and neurulation (fig. 4a). Mirroring this timeseries, the high frequency events depicted by yellow blobs at the top of the heatmap represent periods of synchronized division across the embryo. The dark band in the middle separating two large regions depicts a short transition phase delimiting two phases of ascidian early development. The timing of these stages as reflected in the scalogram matches the timeline of gastrulation and neurulation. Within both phases, the concentric gradients from red to yellow culminating in dense yellow spots in the center of both regions portray increasing morphodynamics.

**Figure 4.**
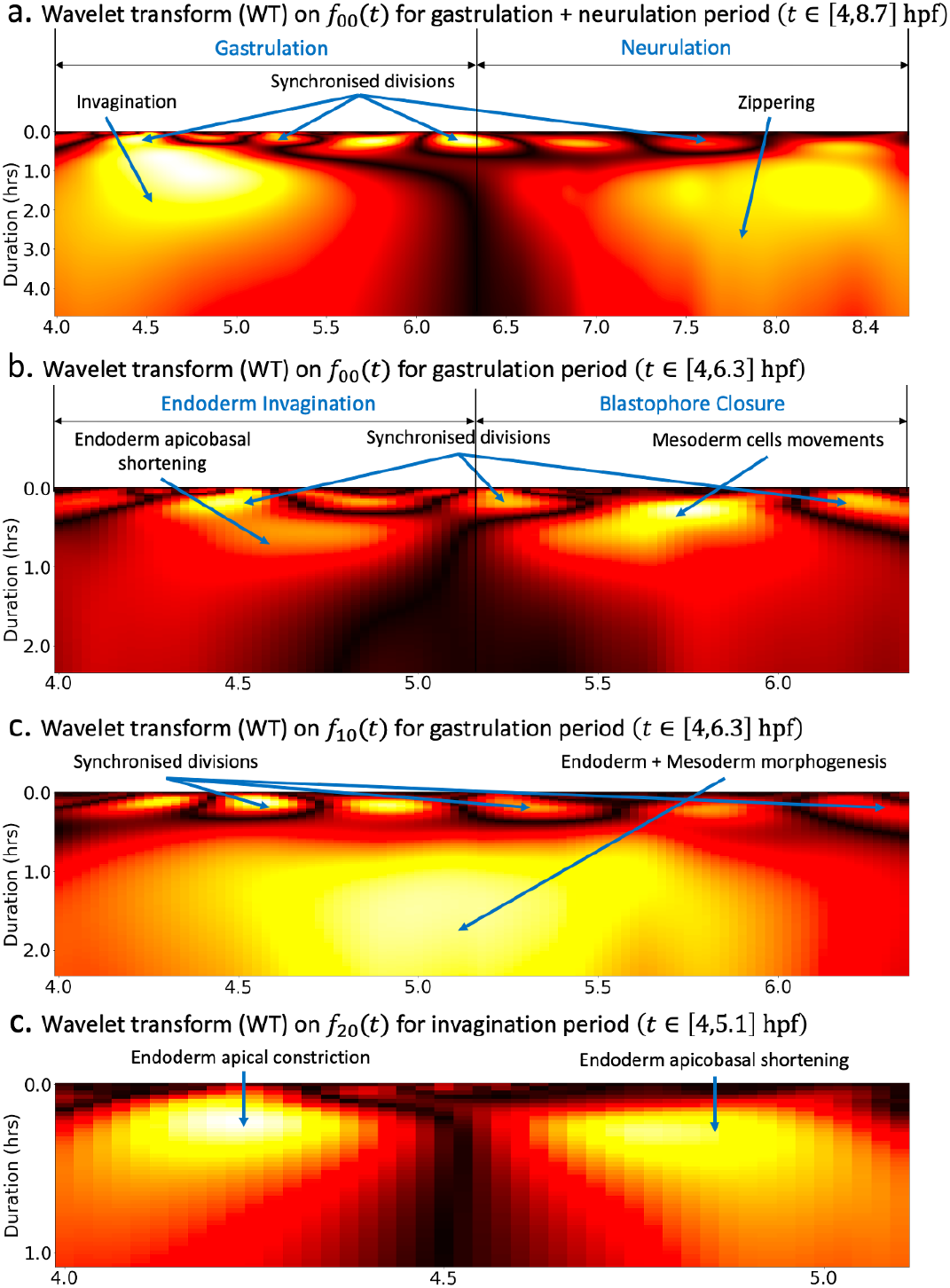
Wavelet analysis highlights multi-timescale modes of morphogenesis. **a)** Scalogram resulting from the ricker wavelet transform applied to *f*_00_(*t*) over the whole period covered by the dataset *t* ∈ [4, 8.6] hpf. **b)** Scalogram resulting from the ricker wavelet transform applied to *f*_00_(*t*) restricted to the gastrulation period *t* ∈ [4, 6.3] hpf. The high frequency events highlighted hererepresent time points of synchronized division across the embryo. The dark band in the middle separating two large red regions indicates that there are two phases of invagination characterized by large deformations and a relatively calm transition phase in between. **c)** Scalogram resulting from the ricker wavelet transform applied to *f*_10_(*t*) restricted to the gastrulation period *t* ∈ [4, 6.3] hpf. Similar to **b)**, the high frequency events indicate synchronized division in the embryo. **d)** Scalogram resulting from the ricker wavelet transform applied to *f*_20_(*t*) restricted to endoderm invagination *t* ∈ [4, 5.1] hpf.

To better understand the specifics of ascidian gastrulation, we restricted the wavelet transform to the gastrulation period (*t* ∈ [4, 6.3] hpf). The resulting scalogram (fig. 4b) shows that ascidian gastrulation unfolds itself in two major phases, delineated on the scalogram by the dark region at the center of the heatmap. The timeline of these events, strengthened by an analysis of topological holes in the embryo (*supplementary* fig. 4a) support the hypothesis that these phases correspond to endoderm invagination followed by the near-closing of the future gut, a process initiated by the collective motion of lateral mesoderm cells known as *blastophore closure*. Both the timeseries (mostly in the negative spectrum) and the scalogram of *f*_10_(*t*) (fig. 4c) adds another layer of validity to this conclusion: the large yellow blob occupying the majority the plot surface highlights that fact that regions of the embryo covered by spherical harmonic *Y*_10_, hence endoderm and mesoderm cells, are subject to intense and prolonged morphological processes.

The first of these two phases, namely endoderm invagination, has been thoroughly investigated in literature. Most notably, it was identified that endoderm invagination was driven by two distinct mechanisms of endoderm single cells ***Sherrard et al. (2010***): first, cells constricted apically by reducing the surface area of the apices, flattening the convex vegetal pole of the embryo setting the stage for invagination. This was followed by animal vegetal shortening of their lateral faces, triggering endoderm invagation. The wavelet transform restricted to the period of endoderm invagination applied to *f*_20_(*t*), whose corresponding spherical harmonic function *Y*_20_(*t*) maps more precisely to the endoderm, beautifully captures this two-steps process (fig. 4d). The timing revealed by this scalogram is in accordance with an analysis endoderm cell shape ratios (supplementary fig. 4b).

### Spectral decomposition of morphogenesis in experimentally perturbed embryo

To assess how our framework adapts to different phenotypes, we set out to conduct a spectral decomposition of morphogenesis in an experimental manipulated embryo. In this particular mutant, MEK kinase was inhibited, which resulted in a massive re-specification of vegetal cell fates, and a disruption of endoderm invagination ***Guignard et al. (2020***). We applied to the mutant dataset (fig 5a *top*) each of the steps in our workflow. First an evolving mesh matching the shape of the embryo at every time point was obtained through the level set scheme. Then, a strain rate tensor field was computed over the surface of the embryo throughout development time (fig 5a *bottom*). A spatiotemporal spectral analysis was subsequently conducted using spherical harmonics on the mutant surface and wavelet analysis of the timeseries of the coefficients of the main harmonic modes. In order to meaningfully compare the dynamics of the mutant development against those of the wild-type embryo, the analysis was carried out at the 64-cell stage.

**Figure 5.**
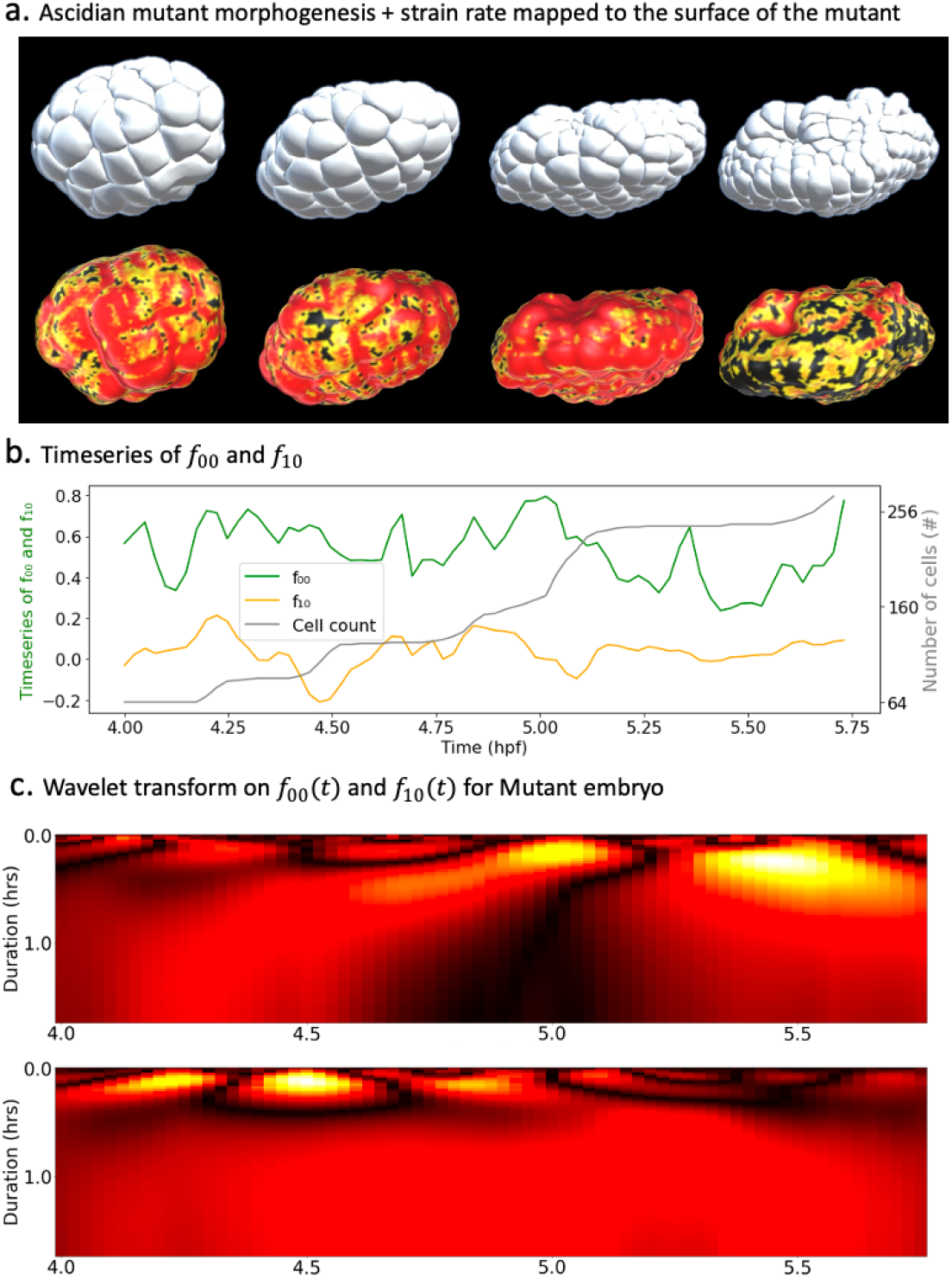
Spectral decomposition of morphogenesis in mutant embryo. **a)** *Top* Ascidian mutant morphogenesis. *Bottom* Spatiotemporal scalar strain rate field mapped to the mutant surface. **b)** Time evolution of the coefficients *f*_00_ and *f*_10_ associated with spherical harmonics (*Y*_00_ and *Y*_10_). The cell population dynamic is also included in the plot for clarity. **c)** Wavelet transform applied on *f*_00_(*t*) (*top*) and *f*_10_(*t*) (*bottom*).

Similar to the wildtype (WT) embyro, the main harmonic modes in the mutant development were *Y*_00_ and *Y*_10_ with respective variance ratios 73.68% and 1.65%, making the timeseries *f*_00_(*t*) and *f*_10_(*t*) the main focus of our examination. The temporal dynamics of these coefficients already reveal major differences between the two strains of ascidians (fig 5b). On the one hand, the drop in the share of *Y*_10_ is telling of the lower order of morphological activity in the vegetal hemisphere. The difference with WT embryos is even more striking when considering that they are deprived of cell division in their vegetal hemisphere. On the other hand, the peaks and lows of *f*_10_(*t*) which coincide with growth in cell numbers, are not restricted either to the negative or positive domain of the curve. This implies that, contrary to the WT, synchronous cell divisions are not restricted to one hemisphere of the embryo.

The wavelet transform applied to timeseries *f*_00_(*t*) and *f*_10_(*t*) yield scalograms which bring further insights into the mutant morphogenesis (fig 5c). On the one hand, the heatmap of *f*_00_(*t*) hints to two distinct phases of the mutant development during the observed time span (fig 5c *top*). The timing of the second phase seems to coincide with the advent of the major wave of cell divisions in the embryo. On the other hand, plots of the scalogram of *f*_10_(*t*) appears to concur with the hypothesis of very low morphological activity in the vegetal pole. The constant red on this heatmap reflects unchanging levels of morphological activity at the vegetal pole of the embryo and confirms the absence of drastic cell deformations that usually drive invagination. This is in alignment with the perturbation induced by MEK kinase which prevents invagination for happening in the mutant endoderm.

## Discussion

Ray Keller’s roadmap of morphogenesis studies establishes a clear path for understanding the biomechanical processes involved in development ***Keller et al. (2003)***. In his proposed workflow, the first step is to determine when and where cells move. Identifying regions of significant morphological activity in space and time has usually followed a script consisting of observing via a microscope the developing system, formulating a hypothesis of what is happening in the system, and subsequently affirming or refuting the hypothesis using qualitative analysis. This method, which has successfully propelled the field of developmental biology to its current heights, nevertheless has some limitations. Distinct morphogenesis events can overlap both in space and time, rendering eye observation vulnerable to misinterpretation. Second, these methods are not automated, hence do not scale.

In this work, we attempted to develop an alternative approach to probing development in living systems. Our approach takes advantage of the recent boom in the availability of single cell shape tracking data to propose a generic method able to identify interesting defining morphological processes through space and time in developing embryos. The method takes as input data consisting of evolving cell geometries and outputs a series of spatial heatmaps showcasing in the time-frequency domains the most salient traits of morphogenesis in the studied embryo. There is however no requirement for segmented cells: the method can be extended to accommodate microscopy imaging data, a feature available in the code submitted. Our framework presents over the traditional eye test method multiple advantages. First, the workflow is fully automated, providing an unprecedented hands-off approach in preliminary studies of morphogenesis. Another outstanding advantage of our workflow over traditional methods is that our workflow is able to compress the story of the development, such that, in a single image, one can grasp the essence of morphogenesis in a system of interest. In particular, our method has been able to neatly discriminate between the gastrulation and neurulation phases of ascidian early development, identify a second phase of gastrulation: *blastophore closure* which follows invagination, reconstitute the two-step process of endoderm invagination during the gastrulation phase, while clearly distinguishing between short scale division events and low frequency tissue-wide deformations.

In order to achieve this fit, raw cell shape data underwent a series of transformations including a level-sets driven homeomorphic map of the unit sphere to the developing embryo’s surface, the computation of the strain rate field of embryo deformations through time using successive iterations of this map, a spherical harmonics decomposition of this strain-rate field, and wavelet decomposition of the most significant spherical harmonics time series. Each of these transformations comes with its own challenges, but also delivers new perspectives for the study of living systems. Our level set scheme excels at defining a homeomorphic map between the unit sphere and surface of the embryo. It goes without saying that in order for the deformed sphere to best match the shape of the embryo, a high sampling of points on the unit sphere is required. A compromise is however necessary between this sampling and, on the one hand, the overall spatial resolution of the original dataset, on the other hand, the induced computational complexity. In its current form, the scheme produces approximations of Lagragian particles only under the assumption of small deformations in the embryo. Hence, the sampling rate during microscopy imaging is of critical importance: the shorter intervals between two successive frames of the movie, the more Lagragian-like the particles are expected to behave.

Given the provision of tracked surface particles meshed at every frame in a triangular network, the evaluation of the strain rate field is straightforward, and enables, among others, a unified description of complex cell-level and tissue-level dynamics ***Blanchard et al. (2009***), such as drastic deformations and synchronised divisions. The accuracy of this field is affected, as previously, by both the spatial sampling of material points on the unit sphere and the timely sampling of morphogenesis frames. Despite the richness in its tensor form, a visualisation of the eigenvalue field derived from this tensor field on the surface of the embryo can already highlight significant processes in morphogenesis. The decomposition of this field into spherical harmonics allows a better appreciation of the spatial patterns of morphological activity in the embryo, each harmonic mapping a region of space. Our spherical harmonics decomposition of morphogenesis results in a set of timeseries of coefficients associated with each harmonic, representing, to the best of our acknowledge, the first comprehensive timeseries-based description of morphogenesis. This transformation unleashes the full power of signal processing tools into studies of morphogenesis. The basis of spherical harmonics being infinite, a challenge here is to discriminate between harmonics that significantly contribute to the composed signal and those that do not. Here, this was done by singling-out harmonics which contributed the most to the variance of the strain rate field. Furthermore, the filtration of principal harmonics modes enables the representation of morphogenesis in a significantly compressed form, in comparison to the initial datasets. This lower dimensional representation of morphogenesis can be helpful, among others, in modelling the physical dynamics of the system ***Romeo et al. (2021***). Another challenge is with the interpretability of the harmonics, which is subject to the alignment of the embryos. The datasets used in this paper presented the advantage that their *y* − *axis* was quasi-aligned with their *vegetal* − *animal* axis. For embryos which do not have this property, prior processing to align them will be required. Alternatively, rotationinvariant representations can be used to appropriately interprete the harmonics.

Despite describing canonical interactions in the space of spherical harmonic functions, our spherical timeseries still represent composed signals in time. We use the *Ricker* Wavelet as a mathematical microscope to zoom-in and zoom-out through these signals in order to identify their fundamentals components. This operation results in two-dimensional time-frequency heatmaps that showcase, for each time-series, the footprint of its canonical high and low frequency components, which can be mapped to biological processes. The sum of these tell the story of morphogenesis in the region corresponding to the spherical harmonic. Here, it might also be useful to wisely target windows of time of interest, and to normalise the data such that interesting transitions can be picked up easily by the wavelet transforms. The resulting heatmaps can be fed to analytic workflows such as deep neural networks for further studies. Example scenarios could include variational studies of morphogenesis processes in different wild-type or mutant embryo. Furthermore, the workflow presented in this paper can be applied to the examination of single cell morphological behaviours in development.

## Code and Data availability

The data used to perform these analyses were obtained from the Morphonet platform (morphonet.org), a platform created and maintained by authors Emmanuel Faure and Patrick Lemaire. The software package developed to perform the langragian mesh construction is publicly available through the following github repository: https://github.com/guijoe/lmg.

