## Supplementary text and figures for "Spectral decomposition unlocks ascidian morphogenesis"

### Spectral decomposition unlocks ascidian morphogenesis (Supplementary material)

#### An overview of ascidian early development

Owing to the relative simplicity of their development, ascidians are an interesting model of development for biologists ***Lemaire*** (***2009***). On the one hand, ascidians embryos are optically clear, which makes them suitable for 3D imaging, observation and the analysis of the dynamics of their cell shapes in fixed and live samples ***Lemaire*** (***2009***). On the other hand, ascidians display rapid development. Whereas the zygote to gastrula transition usually requires several days in other organisms, this process is remarkably fast in ascidian embryos, spanning only a few hours (fig 1a). At the onset of gastrulation, a typical embryo is made of about a hundred cells spatially organized into the endoderm, mesoderm and a cellular mass of unspecified fate ***Lemaire*** (***2009***). Gastrulation is the first large scale morphogenetic process in development, involving drastic rearrangements of cells which reorganise to lay grounds for the future body plan. Lewis Wolpert famous quote *"It is not birth, marriage or death, but gastrulation which is truly the most important time in your life"* ***Hopwood*** (***2022***) emphasises the significance of this process in development. Similar to other large scale processes ***Leptin*** (***2005***); ***Godard and Heisenberg*** (***2019***), ascidian gastrulation is characterised by a dynamic interplay between proliferation and tissue mechanics.

The most dominant tissue-wide movement in ascidian gastrulation is the invagination of the endoderm lineage, which fosters the creation of a gut at the vegetal pole of the embryo. A particularity of invagination in this context is its signature two-steps sequence ***Sherrard et al.*** (***2010***); ***Fiuza et al.*** (***2020***). In the first step, single endoderm cells constrict their apical faces, triggering a coordinated movement that results in the flattening of the endoderm epithelia. In the second step, proper invagination occurs as a consequence of the contraction of endoderm cells apicobasal axes and the expansion of their basal faces ***Sherrard et al.*** (***2010***); ***Fiuza et al.*** (***2020***). Figure 1d illustrates these findings by showing a sharp rise followed by an equally acute decline in the contribution of endoderm cells apicobasal axis to the variance in their shape. As they constrict their apical faces, the variance across secondary axes is reduced and transferred to the apicobasal. Furthermore, as they shorten their apicobasal axis, the accumulated variance is distributed across secondary axes. The combination of these two steps results in a relatively swift endoderm invagination ***Sherrard et al.*** (***2010***), as shown by the rapid increase in gut radius (fig. 1c).

Proliferation also registers as a preponderant mechanism in ascidian development. Through multiple rounds of cellular divisions, the cell population increases approximately three-folds during gastrulation ***Nishida*** (***1986***); ***Lemaire*** (***2009***) and ten-folds through neurulation (fig. 1b). The properties of cellular division during ascidian gastrulation depend on the embryonic hemisphere on which they are located. Divisions at the animal pole display more regularity than those occurring at the vegetal pole. On the one hand, they have been found to be mostly synchronised in time ***Nishida*** (***1986***). On the other hand, through successive rounds, cells alternate their division plane by choosing a plane perpendicular to the previous round ***Nishida*** (***1986***). In contrast, fewer

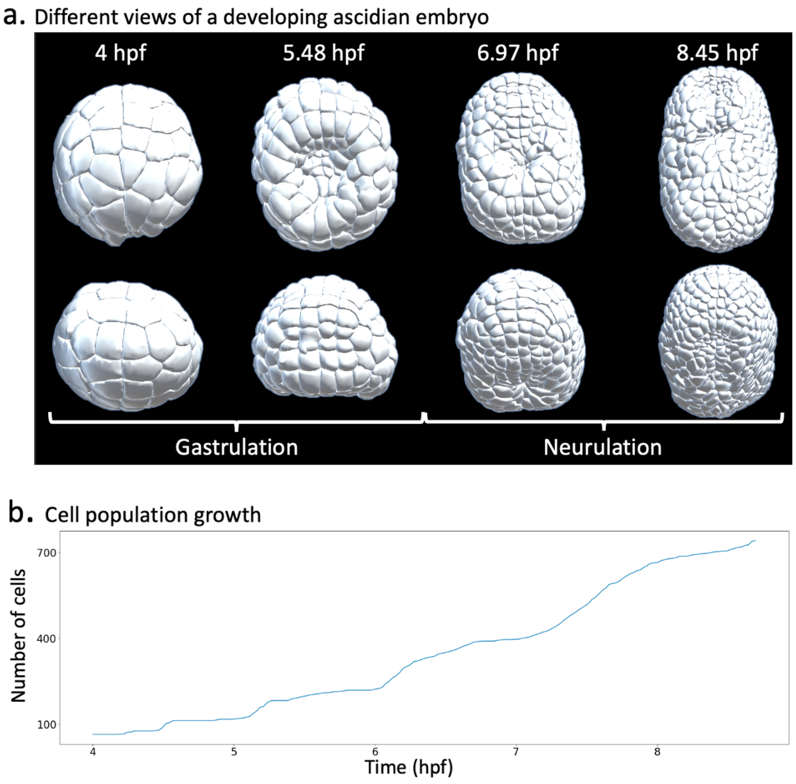


**Figure 1. Ascidian early development Gastrulation.** Tthe early asicidian embryo goes through the phases of gastrulation and neurulation. **a)** Timelines of gastrulation and neurulation in a developing asicidian embryo. **b)** Cell population dynamics.

---

divisions happen at the vegetal pole, and no consistency has been observed in the orientation of divisions ***Nishida*** (***1986***). The question whether these mechanisms influence each other during ascidian gastrulation has also been examined. It was found that embryos in which cellular divisions had been stopped via treatment with noccodazole proceeded without issue to endoderm invagination ***Sherrard et al.*** (***2010***), demonstrating that proliferation is not required for invagination to occur.

As in several species, the development programme in ascidians features neurulation after gastrulation. This important stage consists of transforming the neural plate into the neural tube via the closure of the fold created during gastrulation. Following invagination, cells at the periphery of the gut initiate its closure by inducing a coordinated movement towards the centre. This results in It has been observed that neural tube closure in ascidians evolves in a way similar to the zippering of a hand bag, whereby the gut is closed by the gradual action of cells establishing connections along a linear trajectory ***Hashimoto et al.*** (***2015***); ***Hashimoto and Munro*** (***2018***). The zippering is carried out by the coordinated interactions of about 80 cells ***Hashimoto et al.*** (***2015***), making it a morphological stage with significant mechanical footprint in the storyline of ascidian development.

#### Level set scheme

Here, we propose a method that consists of computing a unique polygonal mesh with fixed vertex networks that evolves to match the shape of the embryo at every single development frame. This method is adapted from ***Zhao et al.*** (***2000***). We consider the following energy, which quantifies the global distance between the two surfaces ($S_1(t)$: real embryonic surface) and ($S_2(t)$: simulated embryonic surface) (eq. 1). This energy reaches a global minimum when the global distance between

$(S_1(t))$ and $(S_2(t))$ goes to zero. Therefore, with sufficiently enough vertices regularly distributed on $(S_2(t))$, both surfaces can be brought to share a similar geometry simply by allowing $(S_2(t))$ points to move along the path of descendant gradients on the energy landscape defined by equation 1.

$$E(S_1(t)) = \left( \int_{S_2(t)} d^2(x, S_1(t)) ds \right)^{(1/2)} \tag{1}$$

Here, $d(x, S_1(t))$ is the distance between a point $x$ of $S_2(t)$ to $S_1(t)$ defined as the minimal distance $d(x, x')$ where $x'$ is a point of $S_1(t)$. $ds$ is the elementary surface area surrounding the vertex with position $x$ on $(S_2(t))$.

For a vertex on $(S_2(t))$ with position $x$, we have the following equation describing its trajectory in the transformation from $(S_2(t))$ to $(S_1(t))$ (eq. 2).

$$\frac{dx}{d\tau} = -\frac{\delta E}{\delta x} \tag{2}$$

Zhao et al. ***Zhao et al.*** (***2000***) show that

$$\frac{\delta E}{\delta x} \approx [\frac{d(x)}{d_{max}}][\nabla d(x) \cdot n + \frac{1}{2} d(x)\kappa]n \tag{3}$$

Here, $d_{max} = \max(d(x)) \approx \left( \int_{S_2^T} d^2(x, S_1^T) ds \right)^{(1/2)}$, $n$ and $\kappa$ are respectively the normal vector and the mean curvature on $(S_0^t)$ at position $x$.

Feeding back equation 3 into 2, we find:

$$\frac{dx}{d\tau} = -[\frac{d(x)}{d_{max}}][\nabla d(x) \cdot n + \frac{1}{2} d(x)\kappa]n \tag{4}$$

Hence, we have the following update rule for a vertex with position $x_i$ in our discrete setting using an explicit Euler scheme.

$$x_i^{\tau+1} = -\left( \nabla_i d \cdot n_i + \frac{1}{2} d_i \kappa \right) n_i \times \Delta\tau + x_i^{\tau} \tag{5}$$

In theory, the scheme such defined is able to bridge the gap between a topological sphere and the embryonic shape. However, the practical implementation of this scheme presents a few challenges.

The first issue we face is that of a proper definition of the surface of the embryo $(S_1(t))$. We recall that our input data can take the form of a discrete set of volumetric cells spatially organised into an embryo, rather than a continuous surface. In this case, we construct the embryonic surface by extracting surface triangles from all external cells and endowing this set with a topology by defining triangle neighbourhoods. For computational efficiency, neighbourhoods are first computed between cells, and then between triangles of neighbouring cells. From this construction, it follows that the distance of a point of space to the surface of the embryo is computed as the minimal distance from this point to all triangles of $(S_1(t))$. In the case that embryonic shape data is in a pixel format, a simple transform of $((S_1^+(t) - S_1(t)))$ - where $S_1^+(t)$ is the dilation operation applied to $S_1(t)$ - can be used to extract the embryonic surface.

The second challenge is the issue of the initialisation of the homeomorphic surface on which the scheme will be applied. For the initial frame $(t = 0)$ of the dataset, in the absence of any reference shape for $(S_1(0))$, $(S_2(0))$ is initialized to a sphere centered at the center of mass of the embryo and whose radius is equal to the maximal radius of the embryo, measured from its center of mass. For subsequent frames $(t)$, in order to maximize computational efficiency and minimize compounding errors arising from the explicit scheme, $(S_2(t))$ is initialized to a weighted average between the minimal sphere circumscribing $(S_1(t))$ and $(S_2^{(}t - 1))$. Other methods of initialisation can be used, including a sparser version of $(S_1^{(}t))$ on which vertices are added.

Inherent to this initialisation problem is the question of the number of vertices required to meaningfully represent the embryonic surface. It is noteworthy here to recall that the surface of the embryo riddled with curved apical faces of cells and discontinuous cellular boundaries, which increase with time: the development time spanned by the dataset features cell population sizes ranging from 64 surface cells all the way up to 562 surface cells. The basic spherical shape is obtained from an icosahedron of 12 vertices whose distances to the center are normalized. Additional vertices are obtained by iterative *butterfly* subdivisions of this spherical mesh ***Hardy and Steeb*** (***2008***). Supplementary figure 2 portrays how well simulated cell junctions (*green points*) mirror the behaviour of actual cell junctions (*red points*) as a function of the number of mesh vertices chosen. As the number of vertices increases, the average relative distance between real and simulated material particles, normalised by the average length length of cell apical faces, decrease (Supp. figure 2b). However, the relative between simulated and real material particles at consecutive frames does not vary much with the number of vertices (Between $5 and 8\%$ Supp. figure 2c). The slight increase observed is most likely due to the need for longer iterations of the gradient descent scheme as the number of vertices grows.

The level set scheme thus described is applied to every frame of development, transforming the initial dataset from a time-series of cellular meshes with arbitrary topologies to a single surface mesh with unique topology, but evolving shape, matching the surface of the embryo at each frame.

#### Strain rate computation

The strain rate tensor field is computed as the symmetric part of the gradient tensor of the velocity field. In order to minimize undesired local artifacts arising e.g. from the inherent errors to experimental measurements and the determination of the embryo surface mesh, the field is smoothed using a Gaussian mask spanning the 1-ring and 2-ring neighbourhood of each vertex. Analytically, the determination of the strain rate tensor field obeys the equations in 6

$$D(t) = \frac{1}{2}\left(L(t) + L^T(t)\right) \tag{6}$$

Here, $L(t) = \nabla v(t)$ and $v(t) = \frac{1}{\Delta t}\left(x(t+\Delta t) - x(t)\right)$, and $x(t)$ is the position of the particle at time $t$.

The determination of the gradient of the velocity field on the mesh entails the determine of different partial derivatives with respect to all spatial dimensions. This is not straightforward in our case of a discrete closed surface embedded in the 3D space. We therefore rely on the work of ***Mancinelli et al.*** (***2018***) on the estimation of gradient fields on triangular meshes to derive expressions for these terms.

In their work, the gradient $\nabla_i f$ of a field $f$ at given vertex $i$ is computed as the mean value over all adjacent triangles $\mathcal{T}(i)$ of the restriction to each adjacent triangle of the field gradient $\nabla_i f_{ijk}$ at $i$. The sum is weighted by the area of each triangle $S_{ijk}$.

$$\nabla_i f = \frac{1}{S_i} \sum_{j,k|ijk\in\mathcal{T}(i)} \nabla_i f_{ijk} S_{ijk} \tag{7}$$

On each adjacent triangle, the field is assumed to be linear and the gradient is evaluated as follows:

$$\nabla_i f_{ijk} = \frac{1}{2S_{ijk}}[(f_j - f_i)n_{ik} + (f_k - f_i)n_{ij}] \tag{8}$$

Here, $n_{ij}$ and $n_{ik}$ represent the normal vectors to edges $ij$ and $ik$ respectively.

#### Spherical harmonics

Spherical harmonics represent a basis of harmonic functions that are useful to decompose signals defined on a sphere. Each of the spherical harmonic functions partitions the sphere into regions

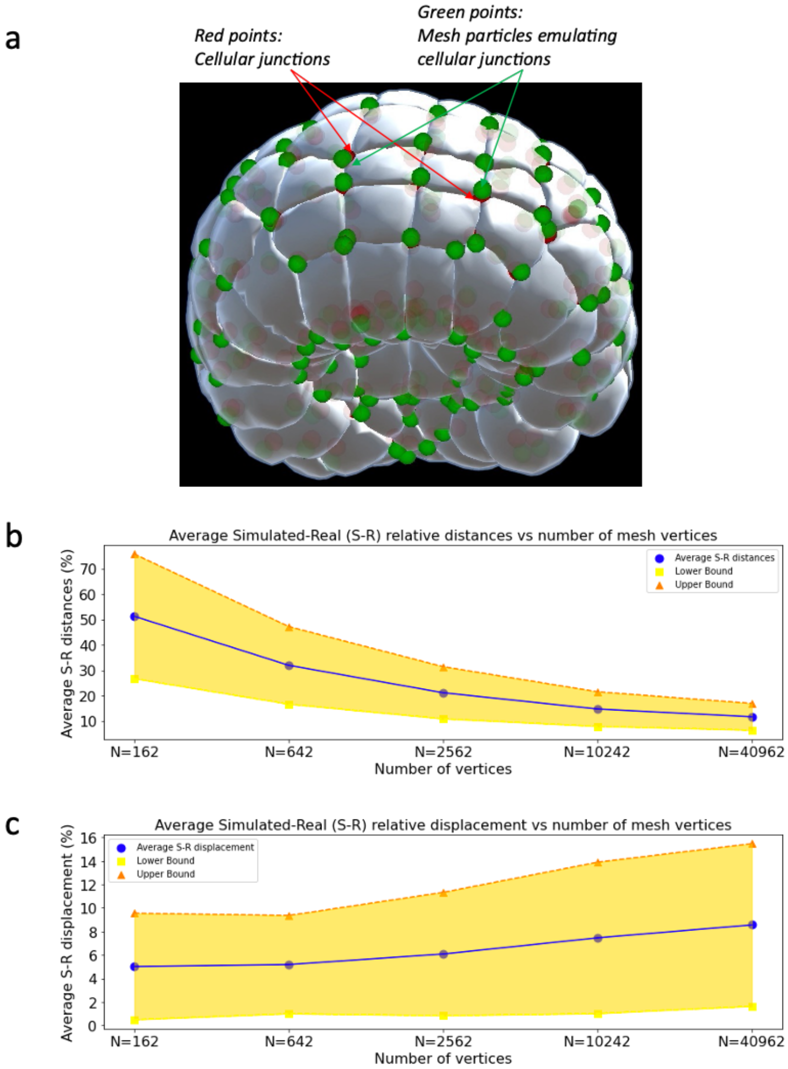


**Figure 2. Level set scheme. a)** Real cell junctions (red) and simulated cell junctions (green) mapped unto the embryo. **b)** Average relative distances between real and simulated cell junctions as a function of number of mesh vertices. The distances are normalised by the average side length of cell apices. **c)** Average relative displacement between real and simulated cell junctions as a function of number of mesh vertices.

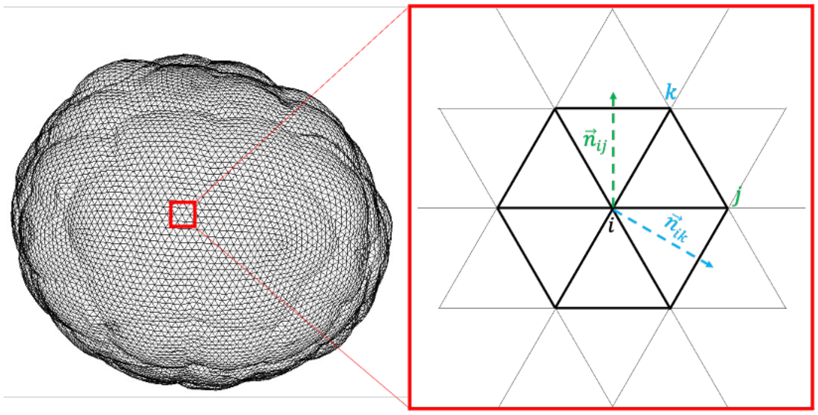

$$\nabla \vec{v}_i = \frac{1}{2 * sum(A_{ijk})} \sum_{(j,k) \in \text{tris}(i)} (v_k - v_i)\vec{n}_{ij} + (v_j - v_i)\vec{n}_{ik}$$

**Figure 3. Discrete gradient computation.**

of space where the signal is positive, negative or equal to zero (Supp. figure 4). Complex spherical harmonic functions are analytically defined as follows:

$$Y_{lm}(\theta, \phi) = \sqrt{(2l+1)\frac{(l-m)!}{(l+m)!}} P_{lm}(\cos\theta) e^{im\phi} \tag{9}$$

Here, $P_{lm}(\mu) = (1-\mu^2)^{m/2} \frac{d^m}{d\mu^m} P_l(\mu)$ and $P_l(\mu) = \frac{1}{2^l l!} \frac{d^l}{d\mu^l}(\mu^2 - 1)^l$

A function $f$ defined on the unit sphere can be expanded using the spherical harmonic basis as follows

$$f(\theta, \phi) = \sum_{l=0}^{+\infty} \sum_{m=-l}^{l} f_{lm} Y_{lm}(\theta, \phi) \tag{10}$$

Here, the coefficients $f_{lm}$ are obtained as $f_{lm} = \oint f(\theta, \phi) Y_{lm}^*(\theta, \phi) dA$

From these coefficients, an approximation of the original signal can be constructed. Although the spherical harmonics form an infinite basis, in practice, we are restricted to expanding a function up to a maximum degree of $L_{max}$. Supplementary figure 4 shows the decomposition and reconstruction of the strain rate scalar field signal using spherical harmonics. It can be visually observed that as $L_{max}$ increases, the reconstructed signal tends to be closer to the scalar strain rate field (*Supplementary* Fig. 4c), hence gradually capturing finer patterns of the scalar strain rate field. In contrast, but none less interesting, lower values of $L_{max}$ tend to better highlight larger spatial patterns of the strain rate field tensor, suitable for the identification principal modes of the studied dynamics, and regions of significant morphological activity (Supp. fig. 4c).

#### Wavelet analysis: The Ricker Wavelet

Like the Fourier transform, wavelets rely on parametrized kernel functions to represent a given signal. These kernel functions are parametrized in such a way that they can be stretched, shrunk and shifted over the domain spanned by the timeseries to adapt to flexible windows in the time-frequency domain. These properties allow wavelets to pick up short-lived effects of high-frequency processes in the signal in narrow bands of time while also capturing background effects of low-frequency processes. Wavelets can however not indefinitely zoom-in or zoom-out through a signal domain as they are constrained by the so-called *'uncertainty principle'*. The multi-resolution of the

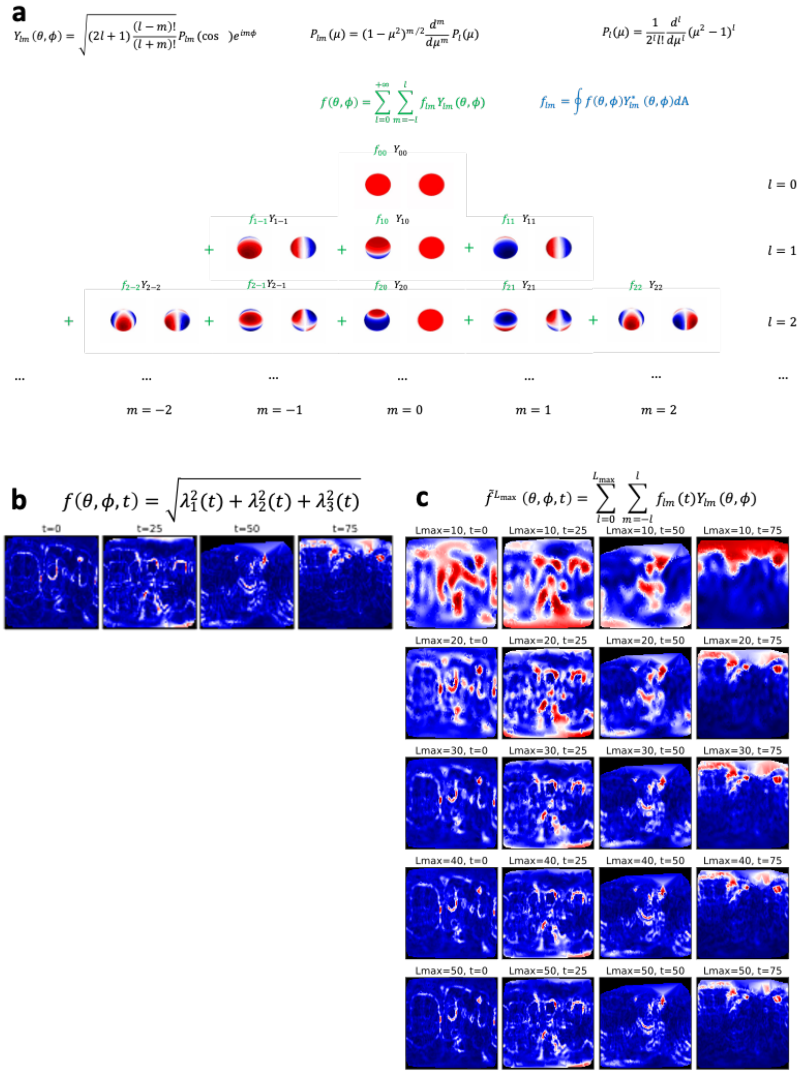

**Figure 4. Spherical harmonics. a)** Analytical expression and spatial representation of spherical harmonics. **b)** 2D representation of the scalar strain rate field. **c)** Reconstructed scalar strain rate field based on the spherical harmonics decomposition up to a degree $L_{max}$

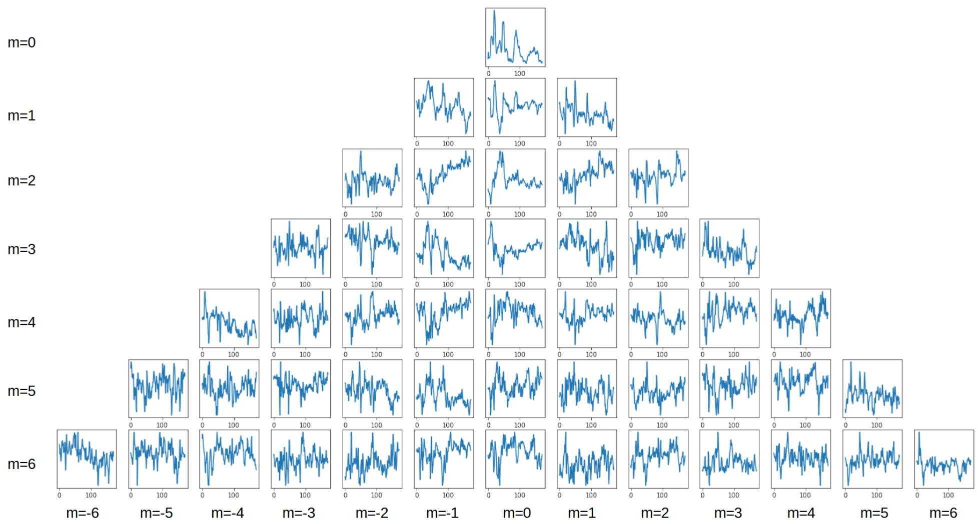

**Figure 5. Spherical harmonics decomposition.** Timeseries of the spherical harmonics coefficients up to l=6.

wavelet transform is hence achieved at the price of a compromise between precision in time and resolution in frequency.

Multiple wavelets kernel exist and are fit for different purposes. In this work, we have made use the Ricker wavelet to analyse the timeseries of spherical harmonic coefficients. The Ricker wavelet can be defined as the negative normalised second derivative of a Gaussian function. Its analytical expression is given as follows:

$$\psi(x) = \frac{2}{\sqrt{3\sigma}\pi^{1/4}}\left(1 - \frac{x^2}{\sigma^2}\right)e^{-\frac{x^2}{2\sigma^2}} \tag{11}$$

Supplementary Figure 6 shows an illustration of the Ricker wavelet transform applied on a composed signal $y(t) = sin(t) + sin(4t)$. Like the Fourier transform, the wavelet can distinguish between the high frequency component ($sin(t)$, tiny yellow blobs at the top of the scalogram) and the low frequency component ($sin(4t)$, large yellow blobs) of the signal. In addition, the wavelet also indicates when the frequencies occur in the signal ($x - axis$), and for how long they last ($y - axis$), thus providing high resolution in both the time and frequency domains.

#### Endoderm dynamics: cell aspect ratios and topological holes

To firmly ground the findings of our spectral decomposition of morphogenesis, we perform a more targeted analysis of the temporal patterns of endoderm morphogenesis. We observe two characteristics of : the length of the apical-basal axis of endoderm cells, and the radius of the gut created by endoderm invagination. Figure 7 present results that perfectly align with the spectral analysis.

In supplementary Fig. 7a, the blue plot represents the average variance of endoderm cells material particle positions across the apicobasal axis during gastrulation. The variations between 4.25 hpf and 5.0 hpf show the two steps of invagination. First, the variance across the AP axis increases rapidly as a result of a decrease in apical surface area. Then, this quantity plummets as a result of apicobasal shortening. The red plot describes the temporal dynamics of endoderm gut radius created by endoderm invagination. The radius peaks around 5.2hpf, which marks the transition between the invagination and the blastophore closure phases of ascidian gastrulation. This radius is computed at every time point using persistent homology, a branch of topological data analysis, which yields itself suitably to the study of topological features such as connected components, voids and holes in point clouds ***Carlsson*** (***2009***).

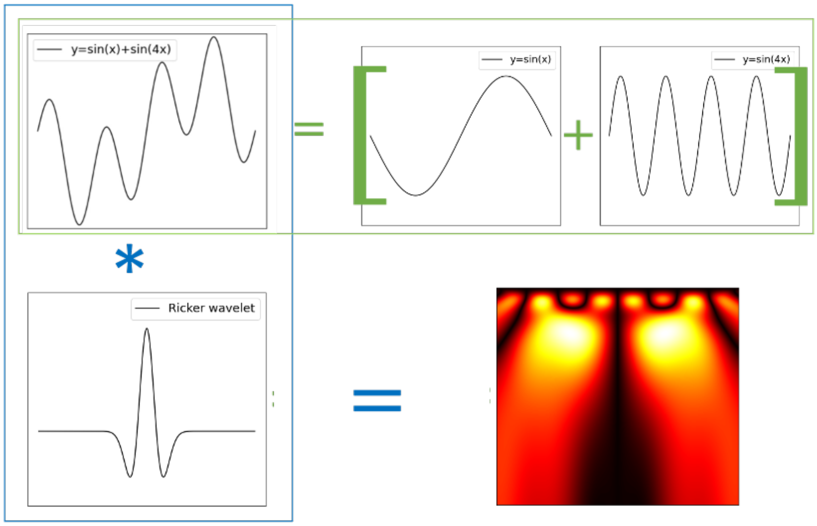


**Figure 6. Wavelet transform.** Ricker Wavelet transform of a composed signal $sin(t) + sin(4t)$. The transform decomposes the signal into its canonical constituents: small yellow blobs for $sin(t)$ and large yellow blobs for $sin(4t)$

Over the entire point cloud consisting of cellular mesh vertices, we sample points from a thin slice of space encompassing material particles of endoderm cells (supplementary fig. 7b). The resulting point cloud is then probed for holes. Before invagination, this region is dense with cellular material particles. However, as invagination progresses, particles leave, creating holes of greater radii between them, which can be captured by persistent homology. The red plot in (supplementary fig. 7a) corresponds to the radius of the maximal hole at each time.

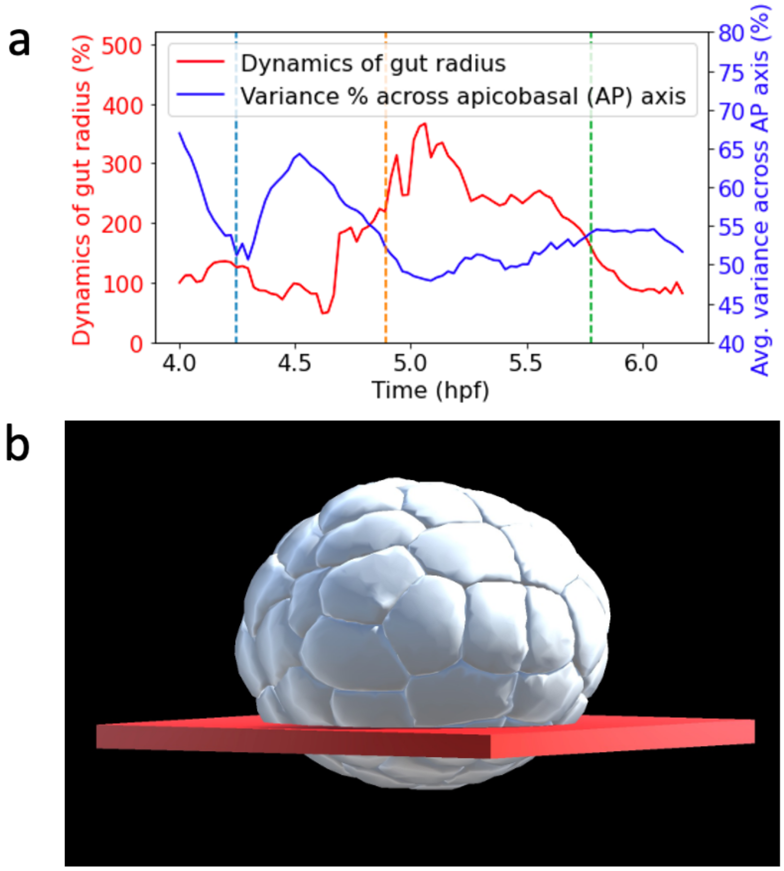


**Figure 7. Endoderm dynamics. a)** Spatial region of point cloud used to observe gut radius dynamics. **b)** Plot of different endoderm dynamics. *Blue plot* Average variance of endoderm cells material particle positions across apicabasal axis during gastrulation. *Red plot* Dynamics of endoderm gut radius.

**Sherrard K**, Robin F, Lemaire P, Munro E. Sequential activation of apical and basolateral contractility drives ascidian endoderm invagination. Current Biology. 2010; 20(17):1499–1510.

**Zhao HK**, Osher S, Merriman B, Kang M. Implicit and nonparametric shape reconstruction from unorganized data using a variational level set method. Computer Vision and Image Understanding. 2000; 80(3):295–314.
